# Orthotopic injection of an established syngeneic mouse oral cancer cell line (MOC1) induces a robust draining lymph node response

**DOI:** 10.1101/2024.01.12.575399

**Authors:** Vikash Kansal, Brendan L.C. Kinney, Nicole C. Schmitt

## Abstract

**Background:** Preclinical models are invaluable for studies on the pathogenesis and treatment of head and neck cancer. In recent years, there has been growing interest in the use of orthotopic syngeneic models, wherein head and neck cancer cell lines are injected into the oral cavity of immunocompetent mice. However, few such orthotopic models have been described in detail. In this brief report, we describe techniques for injection of mouse oral cancer 1 (MOC1) cells into the buccal mucosa and illustrate the tumor growth pattern, lymph node response, and changes in the tumor immune microenvironment over time.

**Methods:** MOC1 cells were injected into the buccal mucosa of C57BL6 mice. Animals were sacrificed at 7, 14, 21, or 27 days. Tumors and lymph nodes were harvested and analyzed for immune cell subsets by flow cytometry.

**Results:** All inoculated mice developed palpable buccal tumors by day 7 and required euthanasia for tumor burden and/or weight loss by day 27. Lymph node mapping showed that these tumors reliably drain to a submandibular lymph node, which enlarges considerably over time. As in MOC1 tumors in the flank, the proportion of intratumoral CD8+ T cells decreased over time, while neutrophilic myeloid cells increased dramatically. However, the pattern and time course of immune changes in the TME were slightly different in the orthotopic buccal model.

**Conclusions:** When used orthotopically in the buccal mucosa, the MOC1 model induces a robust lymph node response and distinct pattern of immune cell infiltration, with peak immune infiltration by day 14.

## INTRODUCTION

The study of head and neck cancer and other solid tumors requires preclinical models that are as life-like as possible. Some of the earliest mouse models for head and neck cancer involved xenografts, wherein human cells are inoculated into immunocompromised mice. With the explosion of immune studies in head and neck cancer, the need for syngeneic, immunocompetent mouse models quickly became apparent. Among these, the mouse oral cancer (MOC) cell lines are among the most popular and well established. These cell lines were established from 7, 12-dimethylbenz(a) anthracene (DMBA)-induced oral squamous cell tumors in mice and have been used in numerous preclinical studies of oral cancer.^1^

In recent years, concerns have arisen about using head and neck cancer cell lines in the subcutaneous flank, which may not represent the immune microenvironment of the upper aerodigestive tract. Not surprisingly, in recent years multiple laboratory groups have transitioned to using orthotopic models, injecting murine cell lines into the buccal mucosa, tongue, or other sites. Although several recent studies have used the MOC models and other head and neck cancer cell lines orthotopically,^2-5^ to our knowledge no studies have provide comprehensive details on how to perform the injections, tumor growth patterns, lymph node responses, or influx of tumor infiltrating lymphocytes in the MOC1 model. Indeed, the evolving tumor immune microenvironment has been well established for MOC1 tumors placed in the flank,^6^ but not in the oral cavity. Prior to using orthotopic MOC1 tumors to study treatment responses, we elected to first study the natural history of these tumors in order to best determine when and how to initiate treatments.

In this brief report, we describe our experience thus far with buccal MOC1 tumors including tumor growth characteristics, lymph node response, and dynamic aspects of the tumor immune microenvironment. This information may be useful to other investigators who may be interested in using the MOC1 model in orthotopic fashion.

## METHODS

### Cell line

MOC1 cells were obtained from Kerafast and maintained as previously described ^7,8^. Cells were regularly tested for Mycoplasma contamination and cultured for no longer than 3 months or 20 passages before use.

### Antibodies and reagents

Fluorescent-conjugated flow cytometry antibodies for mouse tumor experiments were obtained from BD Biosciences, Biolegend, Miltenyi or Abcam (see previously published flow cytometry panels).^9^

### In vivo orthotopic studies

Wildtype, female C57BL/6 mice aged 6-8 weeks were obtained from Taconic. MOC1 cells were prepared as a suspension in sterile PBS with 30% Geltrex and kept on ice. Mice were briefly anesthetized with the isoflurane drop method, then gently restrained by one investigator during injection by another investigator. A total of 5 million cells in 0.1 ml volume were then slowly injected into the left or right buccal submucosa about 1 mm posterior to the anterior commissure. Each mouse was monitored for 1-2 minutes during recovery from anesthesia. After optimizing this technique, we performed lymph node mapping by injecting a small volume of diluted methylene blue into the primary tumor, then exposing the cervical lymph nodes as previously described.^4^

Once satisfied with our injection technique and lymph node mapping, a total of 20 mice were injected with buccal MOC1 cells, then tumors were harvested and weighed at pre-designated time points. Tumors were then processed into single cell suspensions as previously described ^10^ and analyzed by flow cytometry. Tumor draining lymph nodes were also collected and mechanically digested for flow cytometry. All animal procedures were approved by the Institutional Animal Care and Use Committee at Emory University (Protocol #202100008).

### Flow cytometry

Single-cell suspensions from tumors and lymph nodes were rinsed in FACS buffer, then stained with surface antibodies and viability dye (Zombie UV at 1:500 or FVS575 at 1:250) for 30 minutes, followed by additional rinsing. Samples were analyzed on a BD Symphony A3 cytometer, then further analyzed using FlowJo (v10.8.1) software. Gating was performed as previously described.^6,9^ “Fluorescence minus one” controls were tested for each multicolor flow panel.

### Statistical Analysis

Data were analyzed by one-way ANOVA with post-hoc Tukey analysis, using GraphPad Prism software.

## RESULTS

### Tumor injection, growth, and lymph node response

Early on, we recognized the importance of injecting the tumor cells relatively far anterior, just posterior to the oral commissure, and just into the submucosa. If the needle is inserted further, it has a tendency to slide back and over the masseter muscle, producing a tumor that is subcutaneous rather than oral. When left to grow in the anterior submucosa, MOC1 tumors often tend to ulcerate into the mouth, creating a very realistic model (**Figure 1**). A shortage of Matrigel during the COVID-19 pandemic also led us to try diluting our cells in Geltrex, which is more affordable and kept at 4 degrees. We have consistently seen 100% of the MOC1 tumors establish within 7 days using Geltrex, without the inconvenience of thawing a frozen extracellular matrix product. The tumors grow relatively fast (**Figure 2A**), causing difficulty with eating by week 3. Thus, we usually provide moist food and gel at the bottom of the cage for all animals no later than week 3. Weight loss often becomes a problem during week 4 (**Figure 2B**), requiring animal sacrifice. Thus far, we have seen similarly excellent growth with the MOC2 (100,000) and MOC22 (5 million) cell lines when inoculated in similar fashion (not shown).

**Figure 1:**
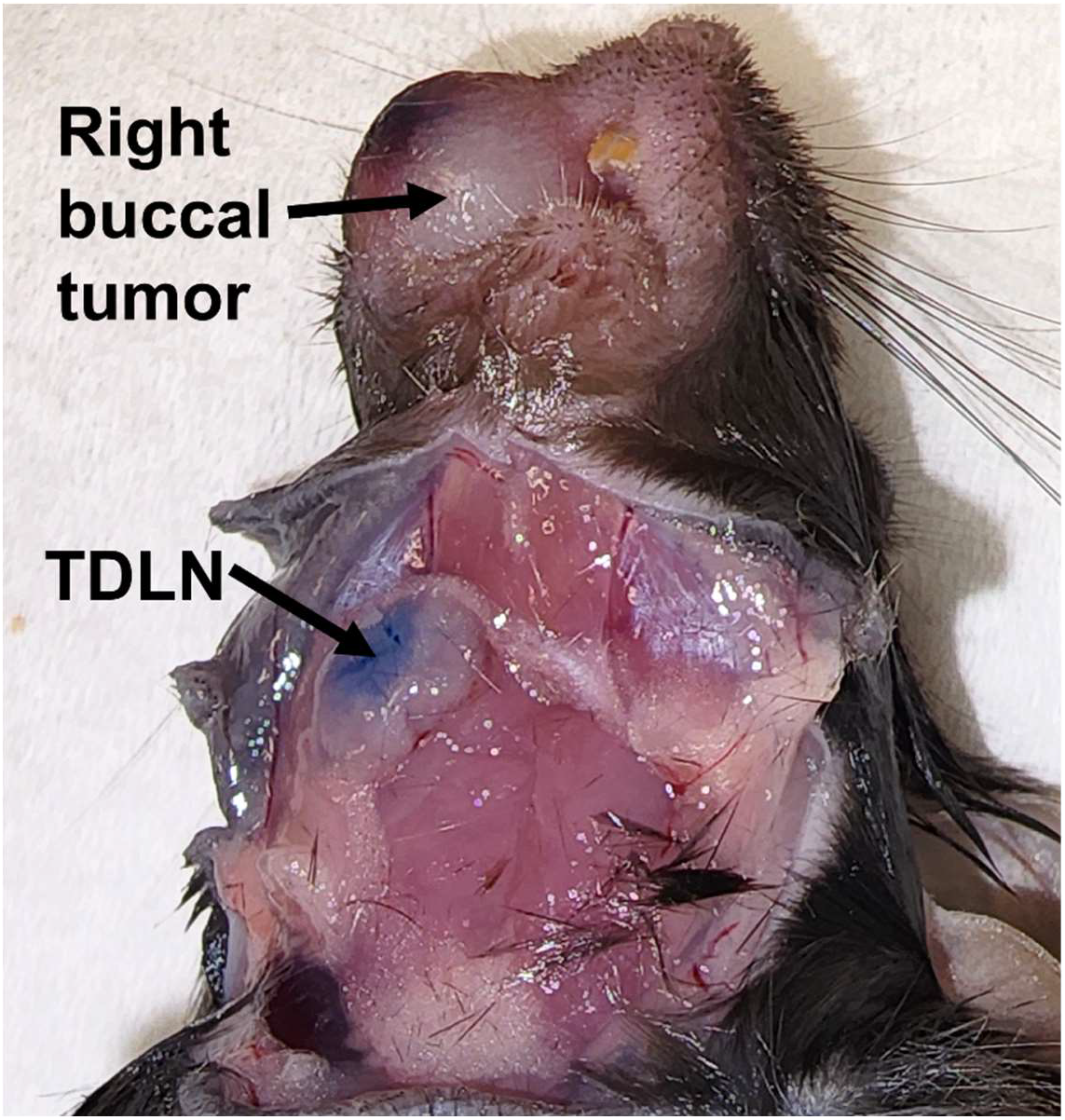
Photograph of buccal MOC1 tumor. The buccal tumor has been injected with methylene blue followed by dissection of the tumor draining lymph node (TDLN), which is adjacent to the submandibular salivary gland.

**Figure 2:**
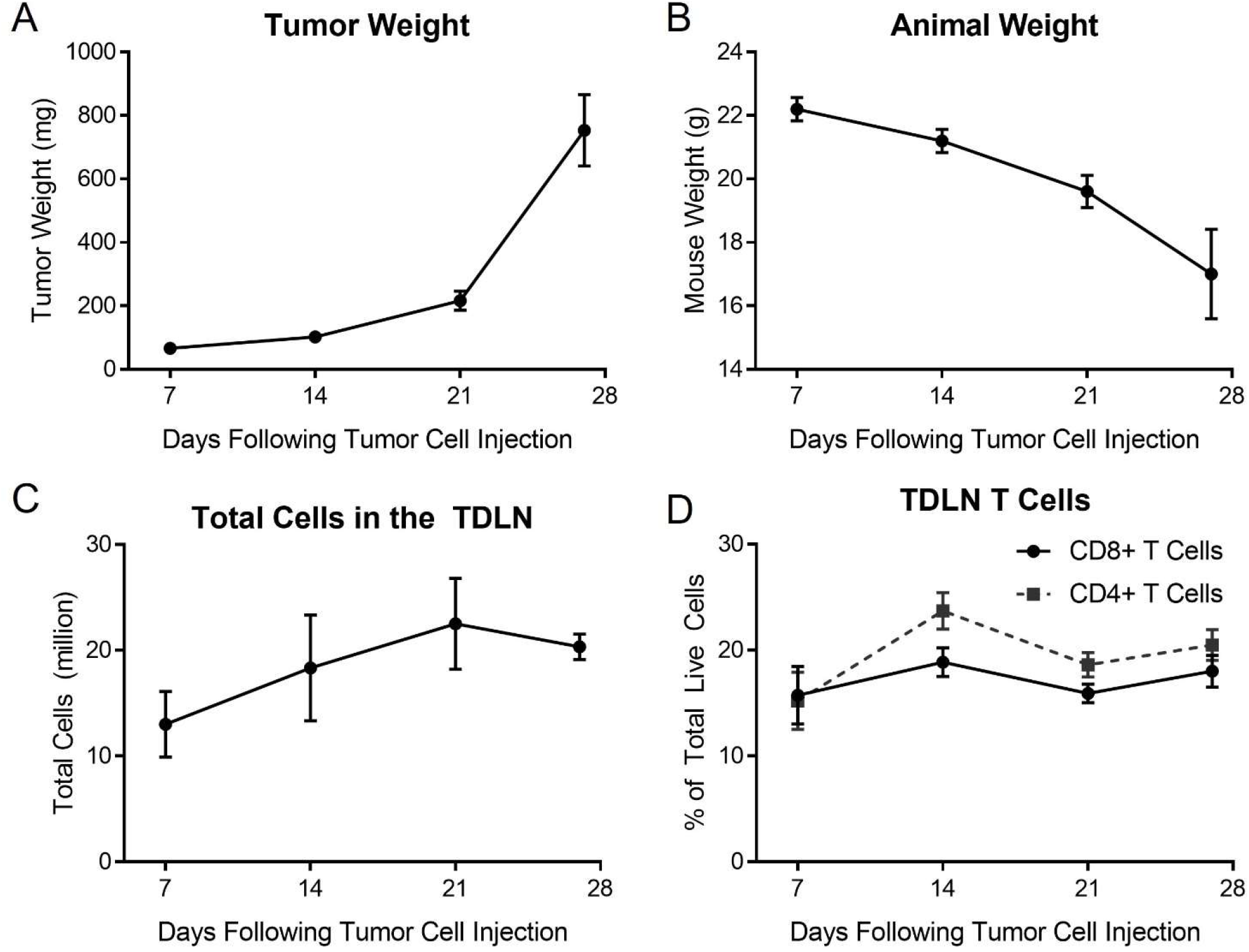
Natural history of the tumor, animal, and tumor draining lymph node (TDLN) after buccal inoculation of MOC1 cells. ***A***, tumor weight in milligrams. ***B***, animal weight in grams. ***C***, total cells collected after mechanical digestion of the TDLN. ***D***, Proportion of CD3+CD4+ and CD3+CD8+ T lymphocytes in the TDLN. Data represent mean ± standard error of the mean.

In our preliminary studies, we also performed lymph node mapping. Diluted methylene blue was injected into the buccal tumors of mice under anesthesia, resulting in a reliable blue sentinel tumor draining lymph node (TDLN) adjacent to the ipsilateral submandibular salivary gland (**Figure 1**), as previously described.^4,5^ We noted the sentinel node to be quite large compared to the TDLN that we typically see when MOC1 is injected into the flank. Indeed, the T cells obtained from the TDLN consistently number in the millions, peaking at day 21, with the proportions of CD4+ and CD8+ T cells staying relatively constant over time (**Figure 2C-D**).

### Changes in the tumor immune microenvironment over time

In subcutaneous MOC1 tumors implanted in the flank, the tumor-infiltrating CD8+ T cells are high at day 10, precipitously dropping while neutrophilic myeloid-derived suppressor cells (MDSCs) increase dramatically; CD4+ T lymphocytes decrease more slowly.^6^ In orthotopic tumors, we found that CD8+ T cells and natural killer (NK) cells were low at day 7, increasing dramatically by day 14, then decreasing just as quickly by day 21 (**Figure 3A**); CD107a, a marker of activation and degranulation on CD8+ T cells, followed a similar pattern, though this did not reach statistical significance (p = 0.067, **Figure 3B**). PD-1 expression on tumor-infiltrating CD8+ T cells peaked by day 21, while its ligand, PD-L1, was highest on tumor cells (EpCAM+) on day 14 (**Figure 3C-D**).

**Figure 3:**
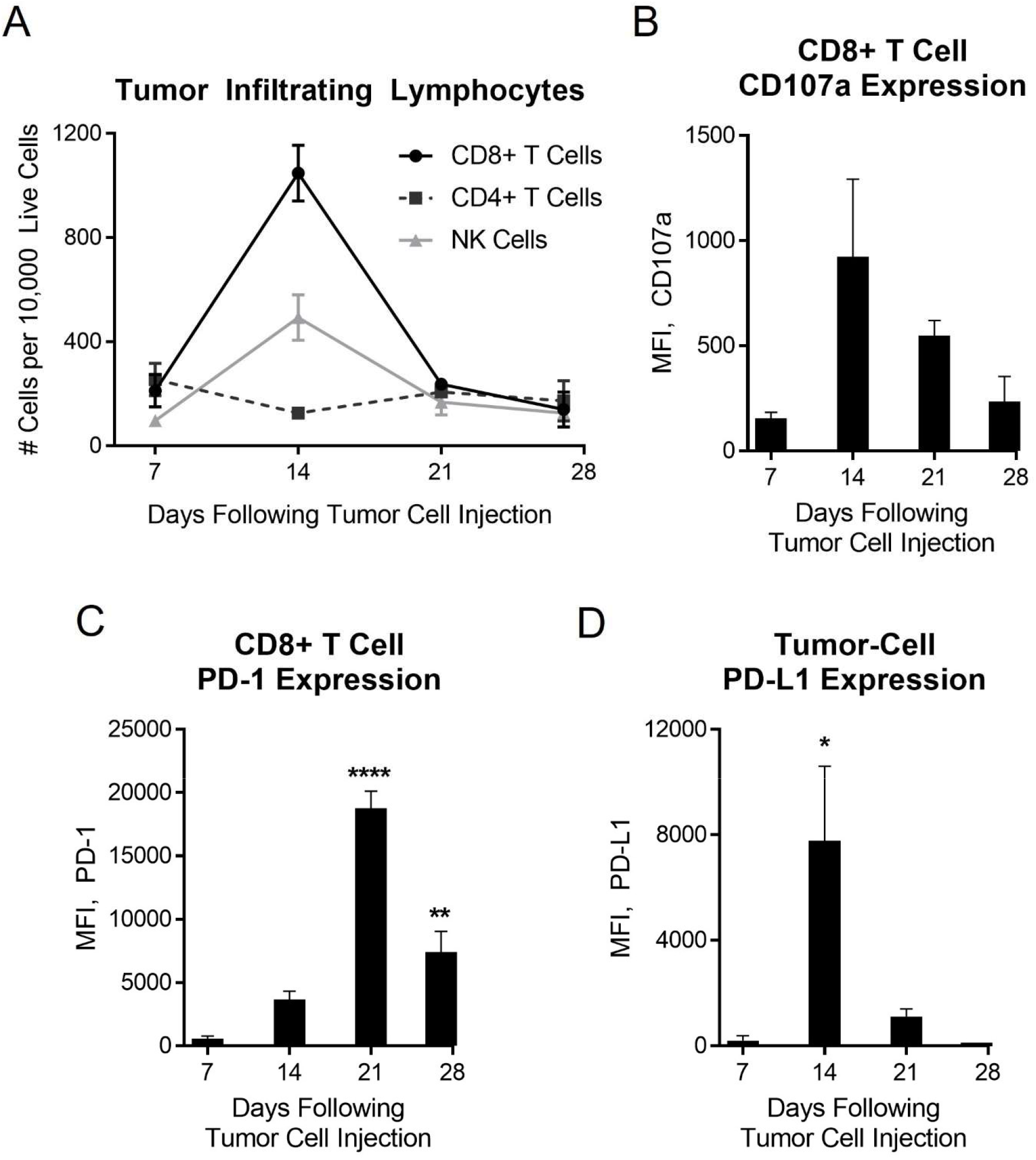
Lymphocytes patterns in the tumor evolved over time. ***A***, Tumor infiltrating lymphocytes (TIL) including CD3+CD8+ T cells, CD3+CD4+ T cells, and CD45+NK1.1+ natural killer (NK) cells. ***B***, surface expression of CD107a on CD8+ TIL. ***C***, surface expression of PD-1 on CD8+ TIL. ***D***, surface PD-L1 expression on CD45-EpCAM+ tumor cells. MFI, mean fluorescence intensity. Data represent mean ± standard error of the mean. *p<0.05, **p<0.01, ****p<0.0001 versus day 7.

Neutrophilic MDSCs (Ly6G-high, Ly6C-intermediate) were relatively constant before skyrocketing on day 27 (**Figure 4A**); by that time, the ratio of CD8+ T cells to neutrophilic MDSCs was at an all-time nadir (**Figure 4B**), similar to what is seen in flank tumors.^6^ Intratumoral macrophage numbers oscillated mildly, while dendritic cells increased modestly over time (**Figure 4C-D**).

**Figure 4:**
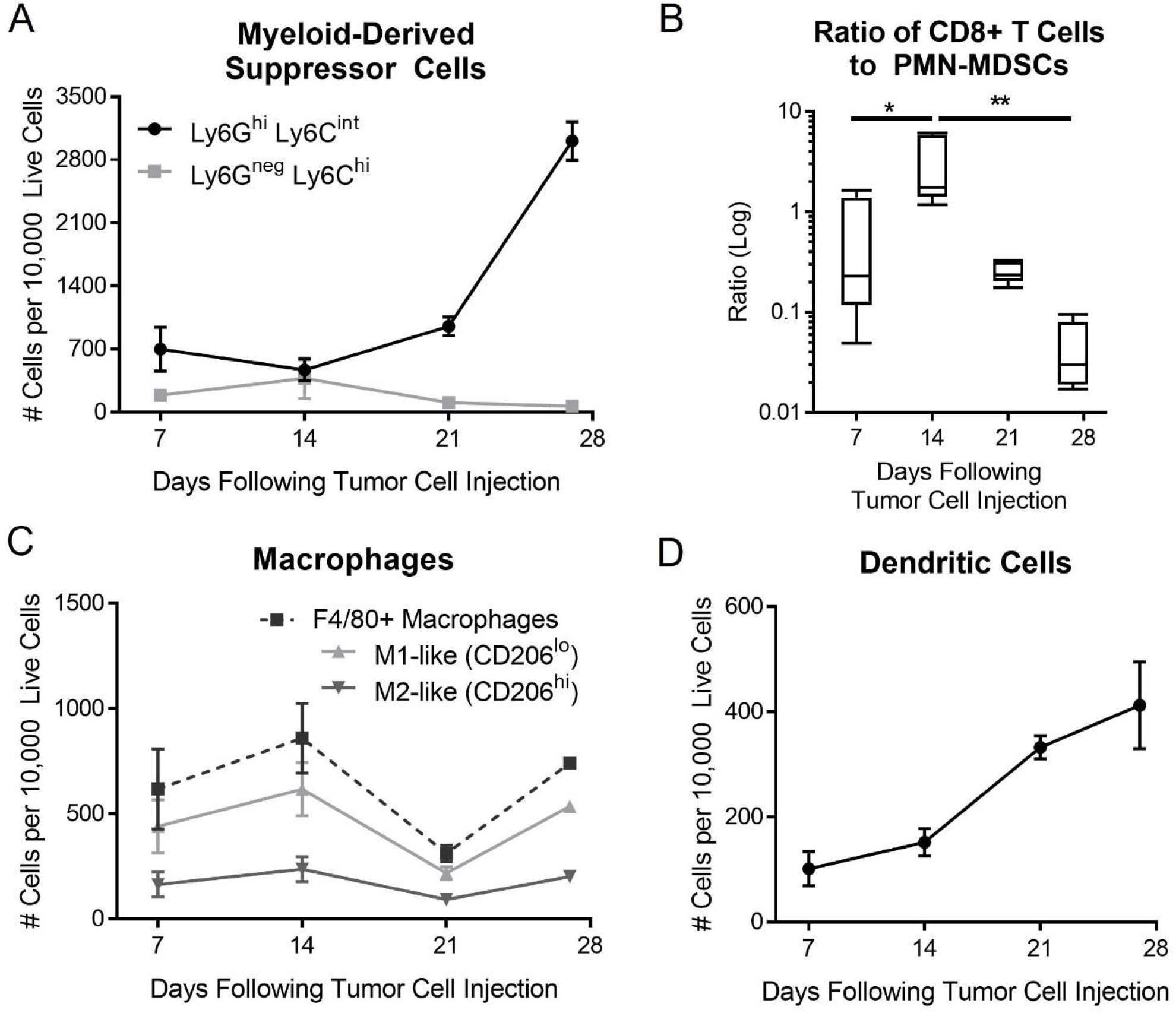
Myeloid cells also shifted over time. ***A***, intratumoral neutrophilic (Ly6G^hi^Ly6C^int^) and monocytic (Ly6G^neg^ Ly6C^hi^) myeloid-derived suppressor cells (MDSCs). ***B***, ratio (log) of intratumoral CD8+ T cells to neutrophilic MDSCs. ***C***, intratumoral CD11b+F4/80+ macrophages, subcategorized as M1-like (CD206^lo^) and M2-like (CD206^hi^). ***D***, intratumoral CD11c+F4/80-dendritic cells. Data represent mean ± standard error of the mean. *p<0.05, **p<0.01.

## DISCUSSION

Lifelike preclinical models are paramount for translational research in head and neck cancer. At the 11^th^ International Conference of the American Head and Neck Society in 2023, a large proportion of the preclinical murine studies presented had used orthotopic models. This is a testament to the interest in such models, despite limited published studies on the methods involved. Rather than allowing each lab to troubleshoot using their preferred murine cell lines in orthotopic fashion, it is helpful to share data on the natural history of these murine models.

Herein, we provide tips on achieving reliable MOC1 tumors in the buccal mucosa, with a robust TDLN response. Similar to previously described patterns for MOC1 tumors in the flank, we see CD8+ T cells decreasing over time while the tumors are heavily infiltrated with neutrophilic MDSCs.^6^ However, the pattern in is slightly different in orthotopic MOC1 tumors, wherein the myeloid cell infiltration occurs more slowly. These differences suggest that immunotherapy may be more effective in the orthotopic model, if started early (i.e., day 7).

Our study has several limitations. Recent studies have used sophisticated techniques such as single-cell RNA sequencing and cytometry by time of flight to study a broad catalog of immune cells in syngeneic mouse models; our study consisted of commonly-studied immune cells delineated by flow cytometry markers. However, a more comprehensive picture of the TME in orthotopic MOC1 tumors is likely to follow in future studies from our lab and others. Our study also used a small number of animals (4-5 per group); however, or subsequent treatment studies (unpublished) have shown consistent results.

In conclusion, orthotopic, immunocompetent models of oral cancer are emerging, and data on the natural history of these models may be useful to investigators who are starting to use them.

## ACKNOWLEDGEMENTS

This study was supported by Winship Cancer Institute and the Department of Otolaryngology at Emory University School of Medicine. This work was supported in part by the Pediatrics/Winship Flow Cytometry Core of the Winship Cancer Institute of Emory University, Children’s Healthcare of Atlanta, and NIH/NCI under award number P30CA138292. The content is solely the responsibility of the authors.

